# Independent Markers of Attentional Shifts

**DOI:** 10.1101/2025.10.31.685868

**Authors:** Kabir Arora, Stefan Van der Stigchel, J. Leon Kenemans, Samson Chota, Surya Gayet

**Affiliations:** Helmholtz Institute, Utrecht University, Heidelberglaan 1, 3584CS, Utrecht, The Netherlands

## Abstract

Visuospatial attention has been extensively studied using a wide variety of markers in the brain and in behaviour. We can broadly classify these markers into two distinct groups: reactive measures, where the presence and strength of attentional biases is inferred from the reaction to a probe stimulus, and proactive measures, which track attentional biases without (or before) the presentation of a probe stimulus. A difference in either type of measure (between two experimental conditions) is typically interpreted to reflect a difference in attentional resources. However, it is not a given that these various reactive and proactive measures of attention all read out the same construct; they may very well tap into independent processes. Here, we measure some of these proactive and reactive signatures of attending a location (changes in cortical excitability as measured by RIFT, alpha lateralization, gaze bias, probe-evoked ERPs, and behavioral reports) in an EEG-eyetracking task (n=23). Though all these metrics indeed correlate with our attentional manipulation, we show that they are not direct correlates of each other, and that of all the proactive measures only the gaze bias predicts subsequent behavioral responses. This demonstrates that existing markers of attention cannot be used interchangeably, and likely capture independent subprocesses of attention.

## Introduction

Visuospatial attention has been extensively studied using a wide variety of markers that reflect whether or not a particular location or stimulus is attended. However, whether these different markers all capture a singular overarching process is still unclear. We can broadly classify these markers of visuospatial attention into two distinct groups: *reactive* measures, where the presence and strength of attentional biases is inferred from the reaction to a probe stimulus, and *proactive* measures, which index attentional biases directly without (or before) a probe stimulus.

Through the reactive approach towards measuring attention, attention to specific locations can indirectly be inferred by presenting a probe stimulus at the location of interest. In the traditional Posner cueing paradigm (Posner, 1980), a specific location is cued, and then a target stimulus is presented at either cued or uncued locations in the visual field. Comparing behavioral performance measures such as reaction time or accuracy (Carrasco et al., 2000; Mangun & Hillyard, 1988; Morgan et al., 1998; Posner, 1980), or the neural response to targets appearing at cued versus uncued locations (Hillyard & Münte, 1984; Luck et al., 2000; Mangun & Hillyard, 1988; Neville & Lawson, 1987; Rugg et al., 1987) provides evidence for the allocation of spatial attention. In this manner the presence and strength of attentional biases is indexed by the (behavioral or neural) reaction to a probe stimulus.

In contrast, the proactive approach attempts to track the allocation of attention itself rather than the consequence it has on the response to a target stimulus. For example, when a particular hemifield of the visual environment is attended, the amplitude of endogenous alpha oscillations decreases in the contralateral hemisphere (Foster et al., 2017; Foxe et al., 1998; Gould et al., 2011; Sauseng et al., 2005). Alternatively, frequency tagging uses Steady State Visual Evoked Potentials (SSVEPs) to obtain a frequency-specific response in the M/EEG signal, the amplitude of which increases when the stimulus is attended (Müller et al., 1998, 2003; Norcia et al., 2015). More recently, Rapid Invisible Frequency Tagging (RIFT) has extended this technique by modulating luminance at frequencies high enough to make the tagging invisible, forming an invisible tracker of spatial attention (Arora, Gayet, et al., 2025; Seijdel et al., 2023; Zhigalov et al., 2019). Finally, the oculomotor system also reflects shifts in attention. Even in the absence of saccadic eye movements, small (<0.5dva) biases in gaze position can reveal the direction and the eccentricity of the attended location (Engbert & Kliegl, 2003; Hafed & Clark, 2002; Van Ede et al., 2019). These measures index the locus of attention without (or before) the presentation of a probe stimulus.

Several measures are used to study attention. However, we also know that attention operates as a multi-level selection process involving the full complexity of the visual processing hierarchy (Serences & Kastner, 2014). Thus, it is not a given that these various measures all read out the same aspect of attention; they likely tap into independent processes that are collectively referred to as attention. In the present study, we measure several proactive and reactive markers of visuospatial attention (RIFT modulations, alpha lateralization, gaze bias, probe-evoked ERPs, and behavioural reports) often used in isolation, concurrently within a single EEG-eyetracking experiment. We show that though all these measures reliably track switches of visuospatial attention, they are largely uncorrelated. This implies that although the results obtained from all of these measures individually are often interpreted to reflect “attention” as an overarching psychological construct, they are likely to capture subprocesses of attention largely independent of one-another.

## Methods

### Participants and Protocol

We recruited 24 healthy healthy human participants (age: 21.95 *±* 1.71, mean *±* std; assigned sex at birth: 18 female, 6 male). One participant was excluded for poor fixation quality, and we thus analysed 23 participants. This sample size was selected in accordance with previous studies studying visual attention using RIFT, alpha oscillations, and gaze biases (Arora, Gayet, et al., 2025; Doherty et al., 2005; Van Ede et al., 2019). Participants reported no history of epilepsy or psychiatric diagnosis, and had normal or corrected-to-normal vision. The study was carried out in accordance with the protocol approved by the Faculty of Social and behavioral Sciences Ethics Committee of Utrecht University, in the labs of the Division of Experimental Psychology, Utrecht University.

Participants underwent a 2-h experimental session. They received procedural information prior to the session. At the beginning of the session, they provided informed consent, date of birth, assigned sex at birth, and dominant hand information. After completion of the EEG setup, participants were seated 72cm from the screen with a chinrest. After eye-tracker calibration, the experiment was explained using a visual guide and verbal script. Following these instructions and 6 practice runs, the *∼*50 minute experiment was conducted. Compensation was awarded when applicable either with €20 or an equivalent amount of participation credits as per Utrecht University’s internal participation framework, and the session was ended. Data collection was conducted between the months of October 2024 and January 2025.

### Experimental Design

We conducted a sustained attention task where participants intermittently switched covert attention between two peripheral locations to respond to target stimuli over the course of consecutive 20s runs (**Figure 1A**).

**Figure 1.**
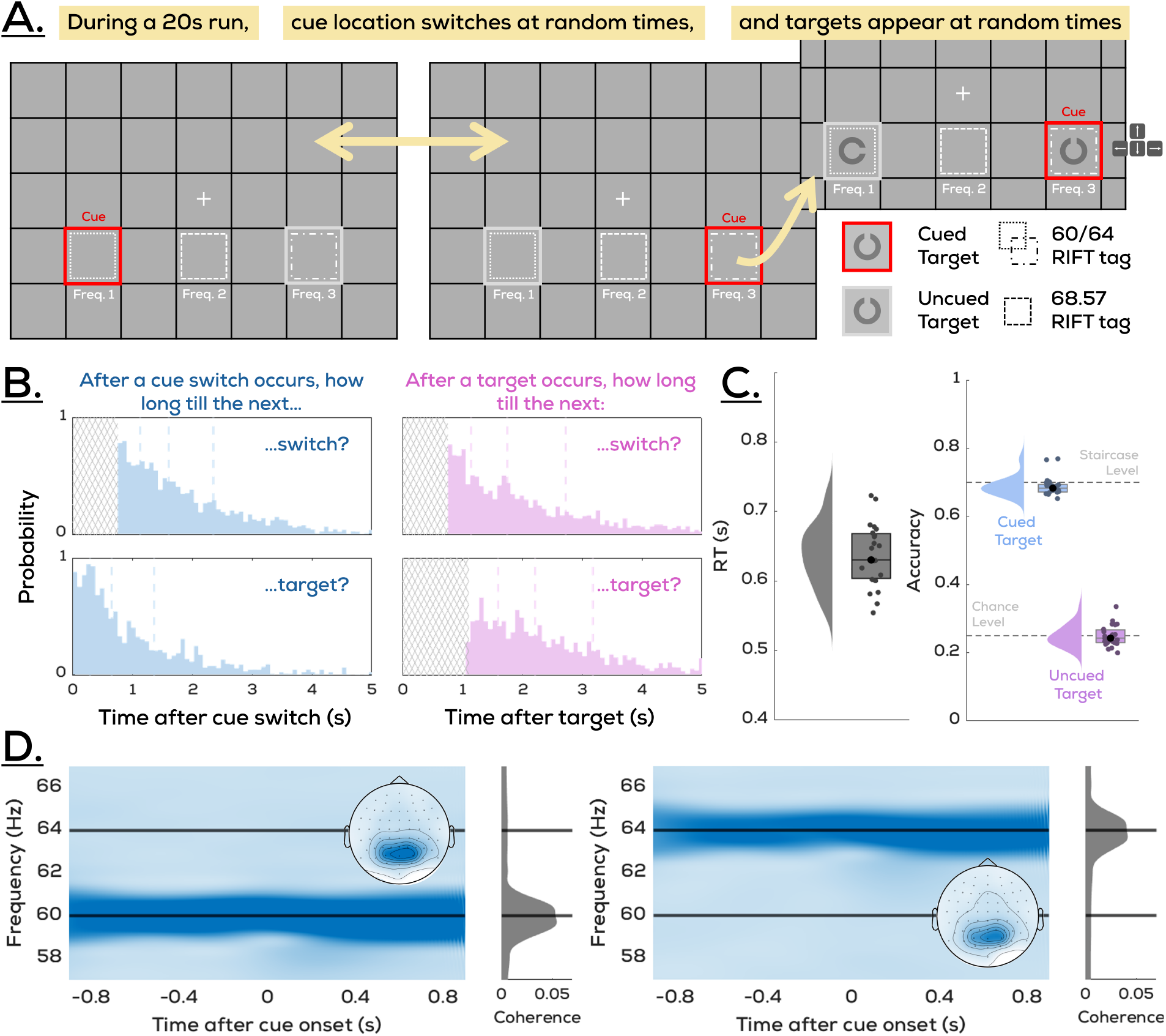
Design and behavioral performance. **A: Task Design.** Participants were shown a grid-like display and maintained fixation in the center. At any given time, one of two lateralized squares was cued, and this cue switched between both squares after random durations. Landolt-C targets would appear simultaneously at both locations at random times, and participants were instructed to report the orientation (direction of the gap) of the target in the currently cued square. **B: Timings**. To prevent participants from being able to predict events (switching of the cue and target onset) we randomised their onsets. However, to simultaneously retain an event-free interval surrounding cue switches for trial segmentation, we placed certain restrictions on the random timings of cue and target onset. Here, the resulting distributions of between task features (cue-cue, cue-target, target-cue, and target-target) are visualized. Dashed vertical lines reflect quartiles. **C: Performance. Left:** Distribution of reaction time for correct responses to the target across participants. **Right:** Accuracy of the reported direction with respect to the cued target, which reflects the staircase value confirming that participants performed the task, and the uncued target, which reflects chance accuracy confirming that the distractor was ignored during report. Shaded regions reflect kernel density estimates of the corresponding participant-wise distribution. **D: Tagging response at 60Hz and 64Hz**. Tagging responses at the two stimulated frequencies were successfully recovered independently from the EEG response (See *Quantifying the RIFT response*).

Each run began with the presentation of a grid layout and a fixation cross at the center of the screen. Participants were instructed to maintain fixation throughout, which was ensured with an eyetracker. Two squares within this grid were highlighted with an outline, their position was one grid-unit below and two units to either the left or right of fixation. One square was outlined red, and the other grey, with frames slightly thicker than the grid lines. The task involved two key events. Firstly, at random moments throughout the run, a landolt-C target would briefly appear (20ms) in each of the outlined squares. Participants were instructed to respond with the orientation of the landolt-C (i.e., the direction of the opening) that appeared in the square with the red outline. The red outline therefore formed a cue that instructed participants which location was relevant at any given time, and we hereafter refer to the square outlined red as the “cued location”, and the square outlined grey as the “uncued location”. Secondly, the cued and uncued outlines switched positions at several random moments throughout the run. This meant that upon each switch (or re-cue), participants had to perform a covert shift of attention to the newly cued location, and maintain it there for unknown duration since the next switch occurred randomly. Importantly, we used RIFT (see *Tagging Manipulation*) to tag three locations continuously throughout the run: the two outlined squares, and a third square directly below fixation forming the ‘middle’ location of the covert shifts. This allowed us to track the allocation of spatial attention by measuring changes in the tagging signal recovered from the EEG response.

We staircased accuracy at 70% using target transparency. This was done to make the task difficult enough that participants could not register the target easily without actually covertly attending the corresponding location. Each 20 second run had 9 or 10 cue switches, and between 5-8 landolt-C target displays. Each participant completed 90 runs (excluding 6 practice runs, not included in analysis) divided into 15 blocks of 6 trials each. Orientations of the landolt-Cs presented in the cued and uncued locations were randomly selected (up, down, left, right) on each trial, meaning that it was possible for both to be the same (with 25% chance).

### Timings

To prevent participants from being able to predict events (switching of the cue and target onset) we randomized their onsets. However, to avoid any ongoing motor activity while a cue occurred, we placed certain restrictions on the random timings of cue and target onset (**Figure 1B**). First, a number of targets (5-8) and a number of cue switches (9-10) were randomly selected for each run. Then, we randomly selected an onset time for each cue switch such that no cues occurred in the first and last 750ms of a run, and no cues occurred within 750ms of each other. Similarly, we randomly selected an onset time for each target such that no targets occurred in the 750ms before a switch, or within 1.1s of each other. Within all permitted windows, timings of cue switches and targets were uniformly random. The intervals between a fixed number of events in a given duration follow an exponential distribution, which, combined with the above restrictions, produced specific distributions for the four types of intervals that occurred in a run (cue-cue, target-target, target-cue, and cue-target).

### Behavioural Reports

Targets appeared in both outlined locations (cued an uncued) simultaneously in the form of landolt-C stimuli with the openings facing either up, left, right, or down. Participants were instructed to respond to the landolt-C in the square currently cued by the red outline, making this the cued target, and the other landolt-C the uncued target. They reported the direction of its opening (up, left, right, or down) with the corresponding arrow key. If a response was not given within 1s of target onset, the trial was marked as incorrect. Participants were informed about this limit prior to the experiment and that they should respond as fast as possible. It was ensured during practice runs that they were achieving this goal, and this is further reflected in the RT results (**Figure 1C**). On average, participants correctly reported the direction of the cued target in accordance with the staircased (70%) accuracy level (accuracy 68.90% ± 2.80%; mean *±* stdev). If the reported orientation matched that of the uncued target more often than chance (25%), this would indicate that participants were not using the cue correctly, however this was not the case (accuracy 24.91% ± 3.11%; mean *±* stdev). Targets to which no response was given in the 1s time limit (mean = 10.4%, median = 9.4% of trials) or an unrelated key was pressed (mean = 0.03%, median = 0% of trials) were labelled as incorrect.

### Stimuli

The screen background was maintained at gray (grayscale: 127.5) throughout the experiment. A black fixation cross (0.4 degrees of visual angle; dva) was present in the center of the screen throughout each trial. The grid consisted of lines (thickness = 0.08 dva) spaced 3 dva apart horizontally and vertically. The frames indicating the cued and uncued locations were twice as thick as the grid lines (thickness = 0.16 dva). At the beginning of the experiment, landolt-C targets (diameter = 2.1 dva) were black with full opacity, but throughout the course of the experiment transparency was modulated through a staircase procedure (Psychtool-Box QUEST algorithm; (Farell & Pelli, 1999), 1999; β = 3.5, Δ = 0.01, γ = 0.5) to maintain accuracy at 70%.

### Tagging Manipulation

We used Rapid Invisible Frequency Tagging (RIFT) to evoke a periodic response from specific locations on the screen (Arora, Gayet, et al., 2025; Seijdel et al., 2023; Zhigalov et al., 2019). Here, this involved sinusoidally varying the luminance at three locations on the screen at specific frequencies. Firstly, the areas within both outlined squares were tagged at either 60Hz or 64Hz. These tagging configurations (60Hz left, 64Hz right; or 64Hz left, 60Hz right) were balanced and randomly assigned to each run. Secondly, the square directly below fixation (“middle location”) was tagged at 68.57Hz on each run (since the middle location was not directly informative about cue-induced shifts of attention, the outcomes of this middle location tag are presented separately in Supplementary Figure S1). Luminance was modulated between white and black, such that the average tag appeared grey and was invisible against the grey (RGB: [127.5, 127.5, 127.5]) background. The edges of the tagged region were smoothly modulated from transparent to opaque to prevent the tagged areas from becoming visible during large eye movements (Minarik et al., 2023). Prior to data collection, the displayed tagging frequency was verified using a BioSemi PhotoCell luminance sensor (BioSemi B.V., Amsterdam, The Netherlands). Temporal precision of the displayed stimuli was continually recorded (at the level of the GPU) during data collection using PsychToolBox’s Screen(‘Flip’) command.

### Display Apparatus

A ProPixx projector (VPixx Technologies Inc., QC Canada; resolution = 960x540px; refresh rate = 480Hz) in a rear-projection format (projected screen size = 48 × 27.2cm) was used for the experimental display. MATLAB and PsychTool-Box (Kleiner et al., 2007) were used for programming the task display.

### EEG Recording and Pre-processing

EEG data were recorded using a 64-channel ActiveTwo BioSemi system (BioSemi B.V., Amsterdam, The Netherlands) at 2048Hz. Two additional electrodes were placed above and on the outer canthus of the left eye respectively. Immediately prior to the experiment, adequate signal quality from all channels was ensured using BioSemi ActiView software. All data analysis was conducted in MATLAB using the Fieldtrip toolbox (Oostenveld et al., 2011). The EEG data were first re-referenced to the average of all channels (excluding poor channels determined by visual inspection, median = 14.1 (mainly frontal) channels, mean = 14 channels). A high-pass filter was applied (0.01Hz), then line noise and its harmonics were removed using a DFT filter (50, 100, 150Hz).

Data were then segmented into trials ranging from 2s before to 2s after cue switches. An ICA was performed to remove oculomotor (blink) artifacts, and trials with other artifacts or noise were removed from further EEG analysis as per visual inspection (median = 5.9%, mean = 7.3%). Baseline correction was performed by averaging (and then subtracting from the signal) a window 0.75s–0.1s before the cue. For ERP analyses, a low-pass filter (30Hz, butterworth filter) was applied.

### Alpha Lateralization

Time-Frequency representations were computed using the ft_freqanalysis function in the Fieldtrip toolbox (Oostenveld et al., 2011). First, spectral analysis was performed on individual trials in the 8–13.5 Hz range (in increments of 0.2Hz). This was done for every channel using 3-cycle Morlet wavelets and baselined (dB) with respect to a window 0.6s–0.3s before cue onset. The 8-13.5Hz frequency range was then averaged to compute alpha power time-series, used to assess the differences in alpha band activity between conditions for each channel. To quantify and visualize the overall lateralization, we computed the difference in average alpha power between trials where the right side versus the left side was being cued. Here, 12 frontal channels were excluded to avoid the influence of consistent gaze position biases (i.e., eye movements artifacts) that we also observed to lateralize depending on cue location. To calculate group-average time-series, differences from the channels that showed a significant group-level effect (negative or positive; bootstrapped 95% CIs excluding 0) were averaged per participant after flipping the sign of channels in the left hemisphere. We also conducted a correlation analysis (See *Correlation Analyses*), relating modulations of alpha power to those of other measures. Since these were conducted trial-wise, they required a different approach. To obtain a single trace on each trial that was more positive if lateralization was stronger, we instead, for each trial, subtracted the mean alpha power of channels showing a significantly negative difference at the group level from that of those showing a significantly positive difference at the group level. We flipped the sign of this difference on trials where the left side was cued.

### Quantifying the RIFT Response

We used periodic stimulation to tag various locations at unique frequencies to track the locus of spatial attention. In order to determine the strength of the tagging response to these RIFT frequencies in the EEG signal, we used two different measures depending on the analysis being conducted. These were coherence and hilbert magnitudes.

### Coherence

For general analyses over all trials, we used magnitude-squared coherence, which is a dimensionless quantity (ranging from 0 to 1) that measures how consistently similar two signals are in both their power and phase. It produces higher values when two trial-wise signals i) oscillate at the same frequency, and ii) maintain the same phase difference across trials (i.e., oscillatory responses across multiple trials are consistently phase-locked). Coherence was computed between a reference wave (pure sinusoids with the corresponding frequency of 60Hz, 64Hz, or 68.57Hz sampled at 2048Hz) and condition-specific sets of trials per channel and participant. When using a reference wave that is equal in frequency and phase on every trial as we do here, this implementation of coherence is qualitatively similar to intertrial coherence. Here, we used the filter-Hilbert Transform (filter-HT) approach to obtain frequency amplitude and phase estimates. Segmented trials were first bandpass filtered (±3Hz) at the frequencies of interest using a two-pass Butterworth filter (4th order, hamming taper). The filtered time-series data were Hilbert transformed. This provided a time-varying instantaneous magnitude (M(t)) and phase (ϕ(t)), the set of which over trials 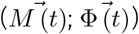 was used to compute coherence as per Equation 1, where x and y subscripts denote EEG data and reference signal features respectively.

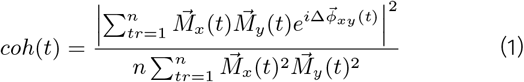

In order to compute coherence spectrograms, coherence was computed for frequencies ranging from 56.8Hz to 67.2Hz in 0.8Hz intervals. In line with previous similar studies, we identified the top 6 channels per individual with the strongest coherence at 60/64Hz across all trials (Arora, Gayet, et al., 2025; Dietz et al., 2025; Hustá et al., 2025; Wang et al., 2025), and any further comparisons across experimental conditions presented here were made using coherence time-traces averaged across these channels. We have, however, previously shown that this exact number of channels selected does not meaningfully impact the strength of the relative differences observed in coherence across experimental conditions (Arora, Gayet, et al., 2025; Arora, Husta, et al., 2025). We baselined the coherence time-traces for each participant using the interval 1.2s to 0.4s before cue onset. This was done separately for each tagging frequency. We obtained overall ‘uncued’ and ‘cued’ RIFT traces by averaging the baselined coherence traces across both configurations (60Hz cued, 64Hz uncued; and 60Hz uncued, 64Hz cued) which were counterbalanced.

### Hilbert magnitudes

For trial-wise correlations between the RIFT response and other metrics, coherence is unsuitable since it considers the phase consistency of a set of trials together. We therefore used a trial-wise measure: Hilbert magnitudes (that is, M(t) as described in Equation 1 above). These values, produced by the filter-HT approach, form the trial-wise, timevarying RIFT response estimates used later for correlation analyses.

### Phase Randomization

We tagged the locations of interest continuously throughout a run, and our event of interest (switching of the cue) occurs at random timepoints during this run. This means that when we epoch the data with respect to target onset, the tagging is not phase-locked across trials. However, the coherence approach that we use to quantify the strength of the RIFT tagging response relies upon phase-consistency. To correct for this, we slightly skewed each trial in time to phase-lock the tagging responses with respect to a given frequency. We thus ran coherence analyses on the data thrice; once with the trial-wise phases aligned to each of the three frequencies tagged (60Hz, 64Hz, 68.57Hz). The maximum temporal skew here was half of one tagging cycle, i.e., *∼*8.3, *∼*7.8, and *∼*7.3ms for 60Hz, 64Hz, and 68.57Hz respectively. This procedure also increases the separability of the tagging responses, since only one frequency is phase-locked at a time in the segmented data. We canthus uniquely track the neural responses to different tagging frequencies in the EEG signal over time (**Figure 1D**).

### Eye-tracking Recording and Analysis

Gaze was tracked using an Eyelink SR (SR Research, Ontario, Canada) eye-tracker. Both eyes were tracked at 500Hz. Immediately prior to the experiment, a 9-point calibration was performed. This calibration was repeated after every 3rd experimental block. The data were segmented into trials of −1.5s before to 2s after cue switches. Blink correction was carried out using custom code adapted from existing work (Hershman et al., 2018). Trials were baseline-corrected with respect to the average position in a 200ms window prior to cue switches. Trials where fixation was not maintained (defined as gaze being further than 2 dva from fixation for more than 50ms of the trial duration) or trials where the baseline was over 2 standard deviations away from the mean baseline position were removed from further eye-tracking analysis (median = 9.8% of trials, mean = 10.8% of trials). A 10ms uniform smoothing filter was applied to the individual position data. To assess the gaze bias at the group level, we averaged gaze position along the horizontal and vertical directions over all trials where the left side was cued and the right side was cued per participant. For trialwise analysis, we used the horizontal component of position over time after flipping the direction of the trials where the left side was cued, such that a positive deviation of gaze position always reflected the cued direction.

### General Statistical Comparisons

Differences in time-varying measures (coherence, alpha lateralization, and gaze position) across conditions were compared over the duration of a trial using a nonparametric cluster-based permutation test (Maris & Oostenveld, 2007). This consisted of four steps. 1) Two traces being compared were subtracted to produce a difference trace. 2) A one-sample t-test was run for each individual time point to detect points on the difference trace significantly different from 0 (p < 0.05). Clusters of consecutively significant timepoints longer than 10ms were identified and their sum of t-values was computed within each cluster to produce a cluster-level t-mass. 3) Then, we randomly flipped the sign of each individual difference trace (hence preserving autocorrelations between timepoints in the null data). We conducted 10,000 repetitions of this process to generate a distribution of expected t-mass values given randomized labels, adding only the largest cluster per repetition. 4) Finally, we checked whether the t-masses of any initially observed clusters were higher than 95% of this distribution. These clusters were accepted as significant. With coherence, we compared the RIFT response evoked from cued vs. uncued item locations first separately for both frequencies, and later averaged across these. For gaze, we tested whether the difference between horizontal gaze position in cued left and cued right trials significantly differed from 0. For alpha lateralization, we similarly tested per channel whether the difference between alpha power in contralateral cued and ipsilateral cued trials significantly differed from 0.

The relationship between target report performance and cue-target intervals was investigated using Inverse Efficiency Scores (IES; reaction time/accuracy). We computed participant-wise IES values for each cue-target interval (100ms bin windows evenly spaced from 50ms to 750ms). We then compared successive cue-target intervals by computing bootstrapped CIs of mean differences. Cue-target interval pairs were labelled significantly different if the 95% CI excluded 0. We observed similar results when using accuracy or RT alone, thus combined both with IES.

### Correlation Analyses

We tested whether the proactive measures of the shift of covert attention correlated with each other, and with the reactive measures using trial-wise, timepoint-by-timepoint correlations. For each proactive measure (lateralization of endogenous alpha oscillations, horizontal bias in gaze position, difference in RIFT response from cued vs. uncued locations), we first obtained trial-wise time-varying values locked to cue onset. Since coherence is computed across a set of trials, we instead used Hilbert magnitudes. We could not directly use the trialwise differences of cued vs. uncued location RIFT responses, since on a given trial cued and uncued locations were tagged at different frequencies (60Hz or 64Hz). Since both configurations (60Hz cued, 64Hz uncued; 64Hz cued, 60Hz uncued) were shown with equal prevalence, we removed the effect of tagging frequency by z-scoring the 60Hz and 64Hz Hilbert magnitude traces. Traces from both frequencies were separately z-scored, but both cued and uncued traces were z-scored together. Lastly, we also compared SNR between the proactive measures qualitatively. For this, using the participant-wise attentional modulation traces of each proactive measure, we computed the mean divided by the standard deviation (M/SD) on a timepoint-by-timepoint basis.

Reactive measures are computed with respect to target rather than cue onset. Thus, we first obtained a value for each reactive measure (RT, and ERP lateralization) following each target. To produce a time-invariant measure similar to RT, ERP lateralizations were then averaged over the time window significant at the group level. We used RT here instead of the IES measure, which includes accuracy as well and was used in other analyses here to assess participant reports. This is because the correlations implemented were trial-wise, and RT offers a continuous measure over trials, but also requires the use of only correct trials, negating the use of accuracy. We excluded trials that were invalid for any of the measures used, i.e., trials where the EEG data had been marked as noisy, where a saccade had been registered as per the eye-tracking data, and where no target was presented following the cue.

Once all trial-wise data was prepared, we ran two sets of correlations. First, we correlated each of the three proactive metrics (RIFT difference, alpha lateralization, gaze bias) with the other two. Then, we correlated the three proactive metrics with the two reactive metrics (RT and target-ERP lateralization). In all cases, we ran both parametric (Pearson coefficient) and non-parametric (Kendall’s Tau coefficient) correlations timepoint-by-timepoint, within each participant. When the proactive (time-varying) metrics were correlated with the reactive (time-averaged) metrics, each timepoint of the proactive metrics was correlated with the time-averaged reactive value. This produced one trace over time per participant of the two correlation coefficients for each pair of measures. We then used these traces to produce 95% bootstrapped CIs over time, and used a cluster permutation test (Maris & Oostenveld, 2007) to determine significance.

## Results

To establish that all the markers we use are sensitive to manipulations of spatial attention, we first show that our reactive measures (target ERP and behavioral perfor-mance) and our proactive measures (alpha lateralization, gaze bias, and RIFT) differ between cued and uncued locations. Then, we test for correlations between the various attentional modulations observed here.

### Reactive measures (target-onset ERP and participant reports) reflect attention, and are correlated

Since a landolt-C stimulus was shown simultaneously at both the cued and uncued locations, any lateralized EEG response following the target onset must be caused by participants using the attentional cue. We obtained a grand average for the difference between contralaterally and ipsilaterally presented cue responses by averaging across all channels with a significant modulation (see **Figure 2A-Top** for example channels). This difference (**Figure 2A-Bottom**) revealed a significant lateralization of the target display evoked ERP around 0.2s (p = 0.0155; cluster t-mass compared to bootstrapped distribution), and from around 0.3s onwards (p = 0.0005). That is, after the presentation of a landolt-C in both cued and uncued areas, the ERP response contralateral to the cued area was significantly lower than the response ipsilateral to the cued area. This is in line with a large body of literature on the N2pc component of attention (Luck & Kappenman, 2012; Stoletniy et al., 2022). A topographical distribution of this right-cued minus left-cued lateralization over a time period of interest as indicated by the channel-level differences (0.14-0.25s) reveals posterior channels with a significant lateralization (**Figure 2B**). In summary, target ERPs in parieto-occipital electrodes indicated lateralized responses after target onset, confirming that the neural response to the target was modulated by attention.

**Figure 2.**
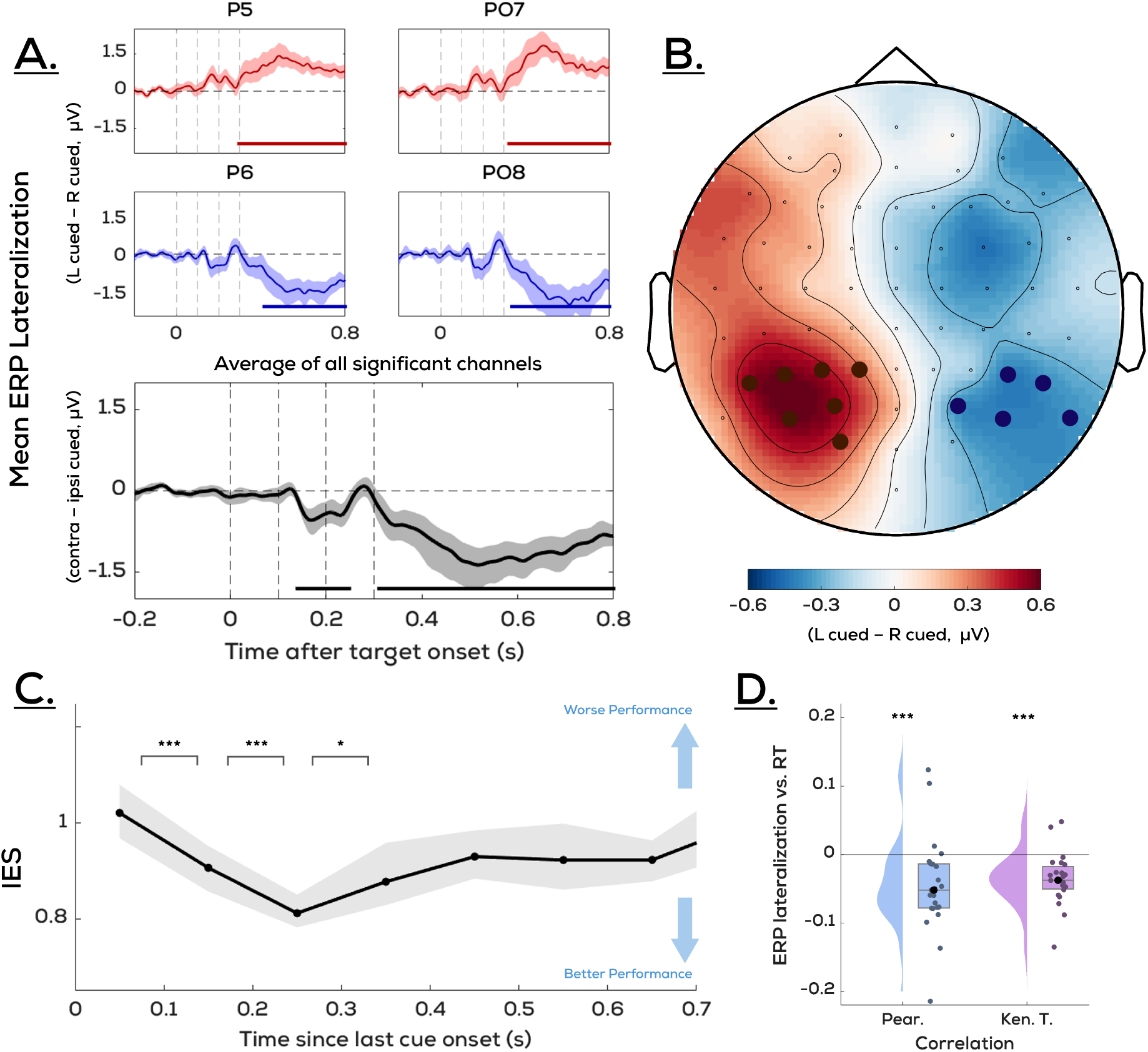
Reactive measures reflect attention and are correlated. A-B: ERPs to targets reflect cued location. Difference in mean target-locked EEG activity between trials where the cue was on the left vs. the right **A-Top:** for selected channels (red = right hemisphere, blue = left hemisphere), **Bottom:** averaged across all significant channels (direction flipped for left channels such that the difference is contralateral minus ipsilateral w.r.t. cue side), and **B:** for all channels averaged across the interval [0.4s, 1s] after cue onset. Shaded regions for all traces represent bootstrapped 95% CIs. Significant channels are represented on the topography with red (positive difference) or blue (negative difference) markers. **C:** performance peaks *∼*200ms after cue. Performance on the target orientation report, measured here using Inverse Efficiency Score (=reaction time/accuracy), binned based cue-target interval. Shaded region represents bootstrapped 95% CIs. Significance labels reflect that the 95% CI of bootstrapped mean differences excludes 0. This statistical comparison was tested only in adjacent bins for clarity. **D:** ERP lateralization correlates with RT. Trial-wise Pearson R and Kendall Tau correlations conducted per participant show a significantly negative correlation, i.e., a stronger target-evoked lateralization links to better performance.

Next, we looked at behavioural reports. In our study, participants were instructed to only respond to targets at the cued location. We therefore cannot compare behavioral responses between the cued and uncued targets. Can we nonetheless obtain a behavioral measure of attention in the present study? We would expect it to take *∼*200ms for attention to shift to the cued location after cue onset. By assessing how behavioral performance evolves with increasing cue-target asynchrony (SOA), we can determine whether participants indeed perform worse on the task in the short moments immediately following cue onset. Behavioral reports can thus be used to test the efficacy of the cue. Here we measure performance by combining accuracy and reaction time (of correct responses) into a joint Inverse Efficiency Score (IES; RT/Accuracy). Indeed, performance increases rapidly over the first few SOA bins (until 200-300ms; 0-100ms vs. 100-200ms: p < 0.0002; 100-200ms vs. 200-300ms: p < 0.0002), after which it slowly starts to decrease again (200-300ms vs. 300-400ms: p = 0.016). This shows that the effect of the cue can be observed in behavioral performance (**Figure 2C**).

### Shift of attention to the cued location is reflected in proactive measures (alpha oscillations, gaze position, and RIFT)

In line with previous literature (Foster et al., 2017; Foxe et al., 1998; Gould et al., 2011; Sauseng et al., 2005), we investigated whether alpha (*∼*10Hz) oscillations decreased contralateral to the hemifield of the cued location. Time-varying differences in alpha power (8-13.5Hz) between right and left cued trials for individual channels (see **Figure 3A-Top** for example channels with a significant modulation) were averaged to present an overall trace (**Figure 3A-Bottom**). Alpha power in response to contralaterally presented cues was lower than in response to ipsilaterally presented cues (p < 0.0005). Visual inspection suggested that this effect is sustained for longer than a second. A topographical distribution of the difference between right-cued and left-cued trials (**Figure 3B**) visualizes the spatial extent of this difference averaged over a time period of interest as indicated by the channel-level differences (0.4-1s). In summary, we observed robust alpha lateralization across channels and time following cue onset.

**Figure 3.**
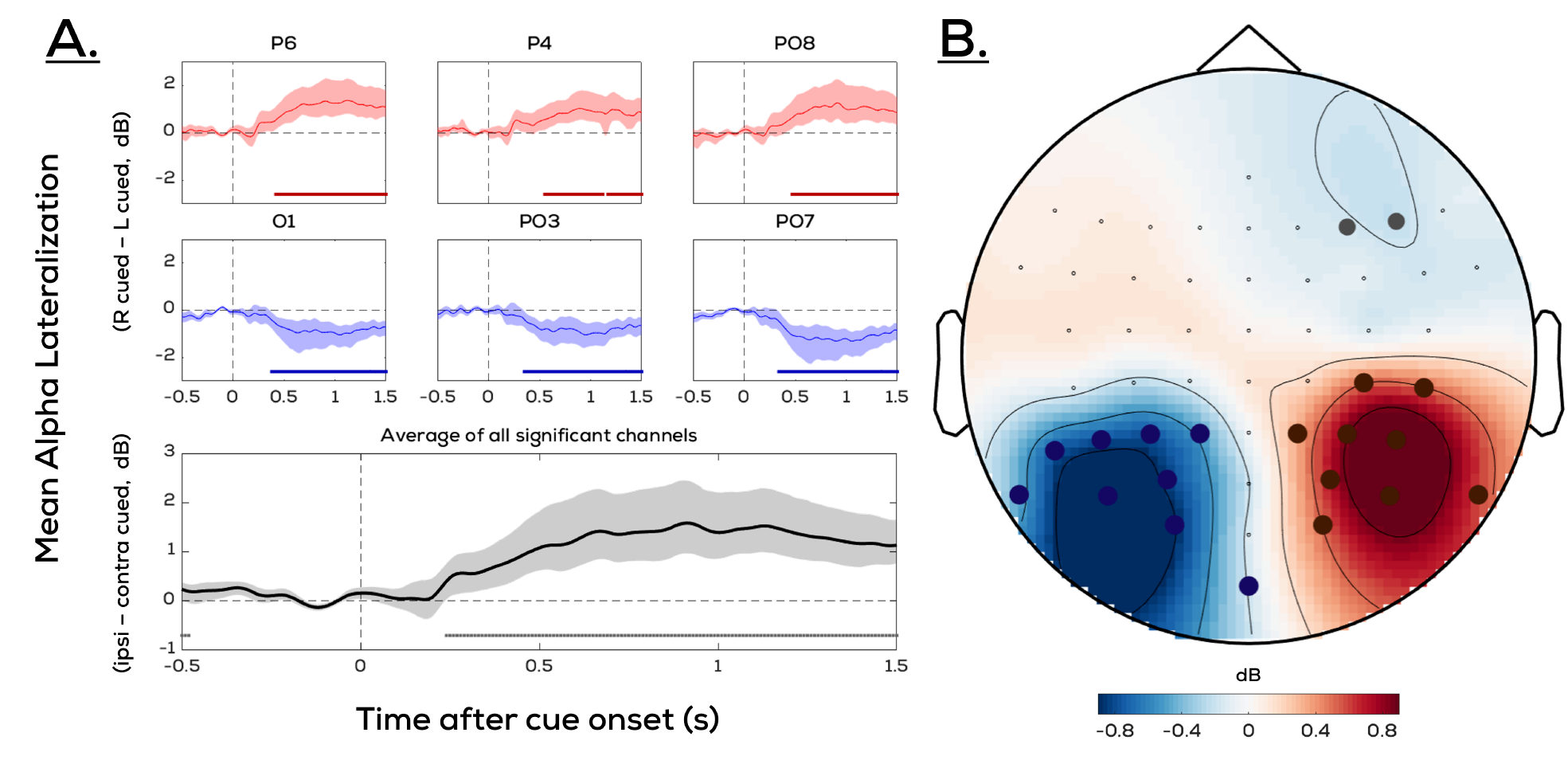
Alpha Oscillations reflect the cued location. Difference in alpha (8-13.5Hz) power between trials where the cue switched to the right vs. to the left **A: Top:** for selected channels (red = right hemisphere, blue = left hemisphere), **Bottom:** averaged across all significant channels (direction flipped for left channels such that the difference is ipsilateral minus contralateral w.r.t. cue side), and **B:** for all channels averaged across the interval [0.4s, 1s] after cue onset. Alpha power is reduced contralateral to the hemisphere corresponding to the cued location. Shaded regions for all traces represent bootstrapped 95% CIs, and bars represent significance based on a cluster permutation test. Significant channels (where the bootstrapped 95% CIs exclude zero) are represented on the topography with red (positive difference) or blue (negative difference) markers.

Even in the absence of saccades, the oculomotor system has been shown to reflect the direction of covert shifts of attention through small (<0.5dva), but systematic, biases in gaze position. First, we tested whether the participant-wise traces of horizontal eye position were significantly biased towards the cued direction for trials where the cue was presented on the right and on the left. This revealed a significant bias of right-cued trials (p = 0.038; cluster t-mass compared to bootstrapped distribution) and an almost significant modulation of left-cued trials (p = 0.05) towards the cued location. Before the cue, a significant modulation of right-cued trials (p = 0.0002) as well as left-cued trials (p = 0.0036) towards the opposite location was seen, likely a result of the previous cue (which was at a random interval, and should therefore yield an effect that is more smeared out in time).

Then, we computed participant-wise difference traces by subtracting left-cued from right-cued traces. These were significantly biased towards the cued location shortly after cue onset (p = 0.036; cluster t-mass compared to boot-strapped distribution) as well as towards the other location prior to cued onset (p = 0.001); cluster t-mass compared to bootstrapped distribution). Again, this flip in the orientation of the effect prior to cue onset is likely a result of the previous cue. Thus, we observed a clear bias in gaze position towards the cued side after cue onset (**Figure 4**).

**Figure 4.**
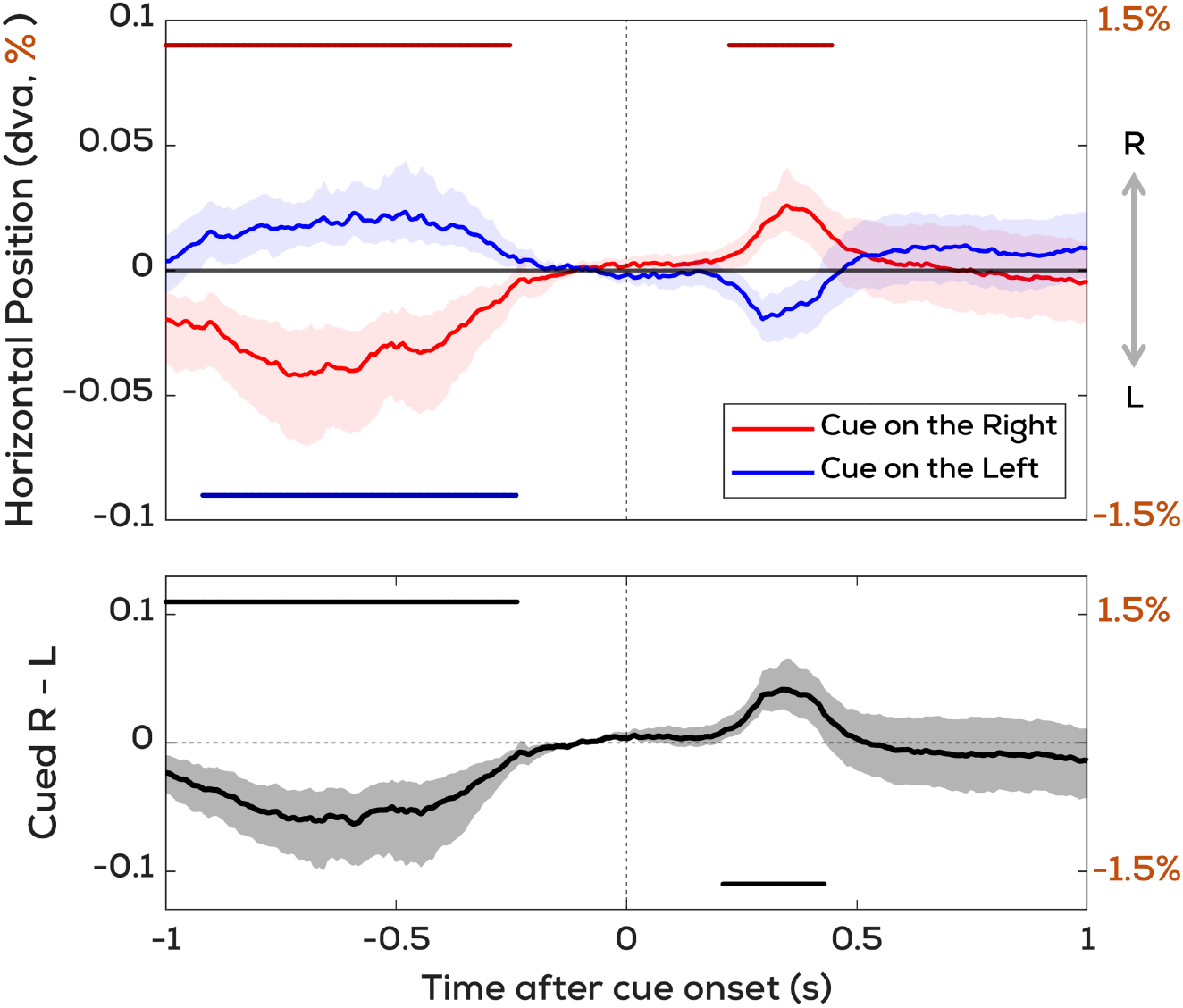
Gaze position reflects the cued location. **Top:** Mean horizontal gaze position for trials where the cue switched to the right vs. to the left. **Bottom:** Difference trace of above. Distances are reflected both in dva and in percentage of the distance from fixation to the center of the cued square. Shaded regions for all traces represent bootstrapped 95% CIs, and significance bars represent significant deviation from chance as determined by a cluster permutation test.

Next, we looked at RIFT responses in the EEG signal. With RIFT, we flicker regions of the screen at specific frequencies (here 60Hz, 64Hz) and the strength of the oscillatory response to these tags offers a measure of covert attention over time. We present ‘cued’ and ‘uncued’ coherence traces (see *Quantifying the RIFT Response* for details) as a measure of the RIFT response from the cued and uncued locations following cue onset (**Figure 5-Top**). We observed two clusters representing a significantly higher RIFT response at the cued location compared to baseline (early, p < 0.0005; and late, p = 0.0015). We also observed a significantly higher RIFT response at the uncued location compared to baseline shortly after cue onset (p = 0.0065), likely a result of a visual transient (cue switch) at this location. Later, a significantly lower response was seen at the uncued location (p = 0.0045).

**Figure 5.**
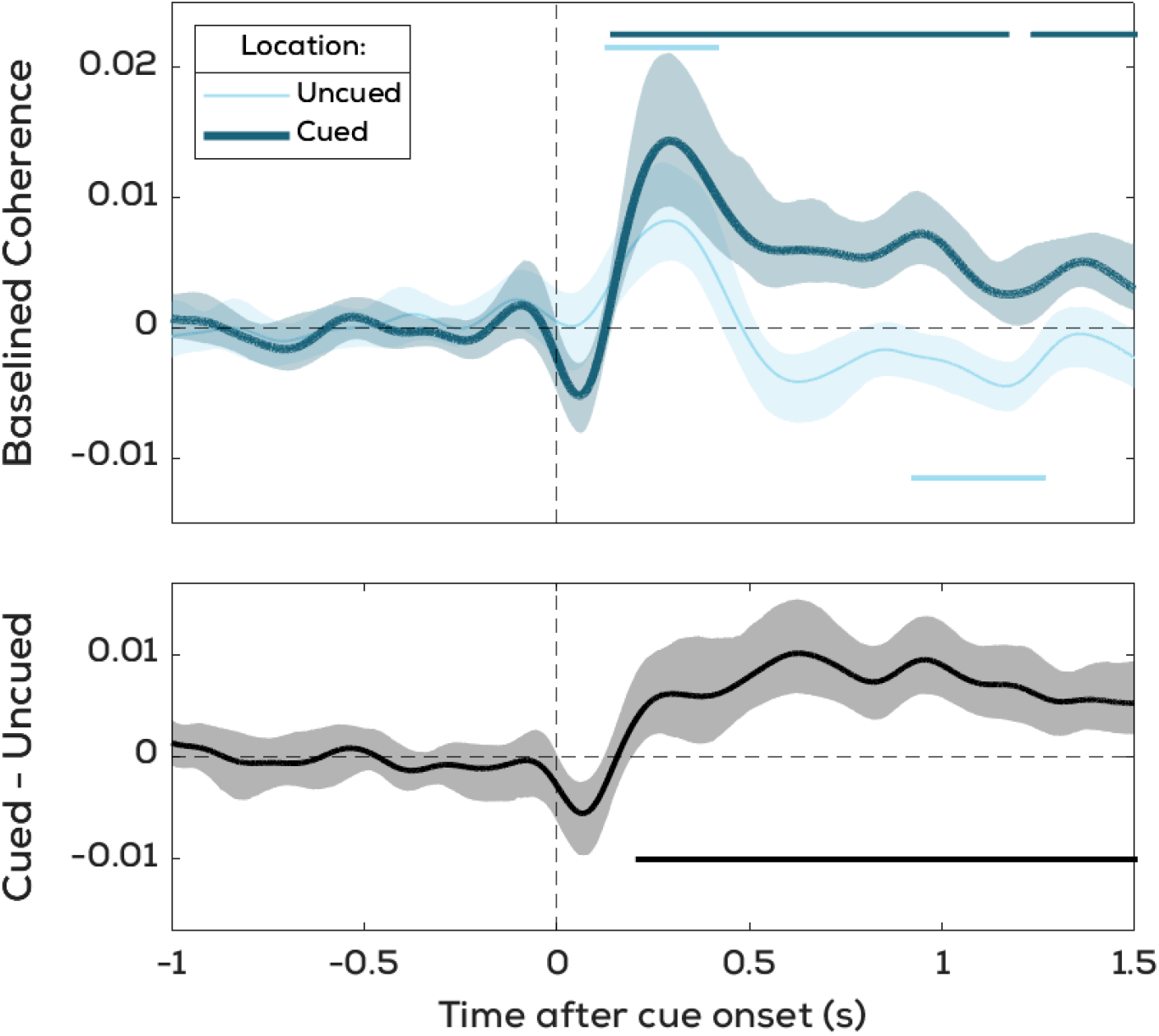
RIFT responses reflect the cued location. **Top:** Mean RIFT responses from the cued and uncued tagging locations. Traces are baselined with respect to the period before cue onset. **Bottom:** Difference trace of above. Shaded regions for all traces represent bootstrapped 95% CIs, and significance bars represent significant deviation from chance as determined by a cluster permutation test.

To assess whether RIFT picked up on the shift of attention to the cued location, we computed the difference trace between the cued and uncued locations (**Figure 5-Bottom**). A significant difference was observed (p = 0.0005; cluster t-mass compared to bootstrapped distribution), reflecting a stronger RIFT response at the cued compared to the uncued location. This confirms that visual processing was boosted at the cued location for a sustained duration following cue onset.

### Proactive measures do not correlate with each other, and only the gaze bias correlates with target reactions

Having established that the spatial cue affected all our metrics (alpha oscillations, gaze position, RIFT, target ERPs, target report), we then tested for correlations between these. We conducted timepoint-by-timepoint parametric (Pearson coefficient) and non-parametric (Kendall Tau coefficient) correlations across trials separately for each participant (see Correlation Analyses section in methods for how data from each metric was prepared). When looking at the proactive measures (gaze position bias, alpha lateralization, RIFT difference between cued and uncued locations), no significant cross-metric correlations were observed (**Figure 6**), tentatively showing that these metrics reflect independent processes of attentional shifts.

**Figure 6.**
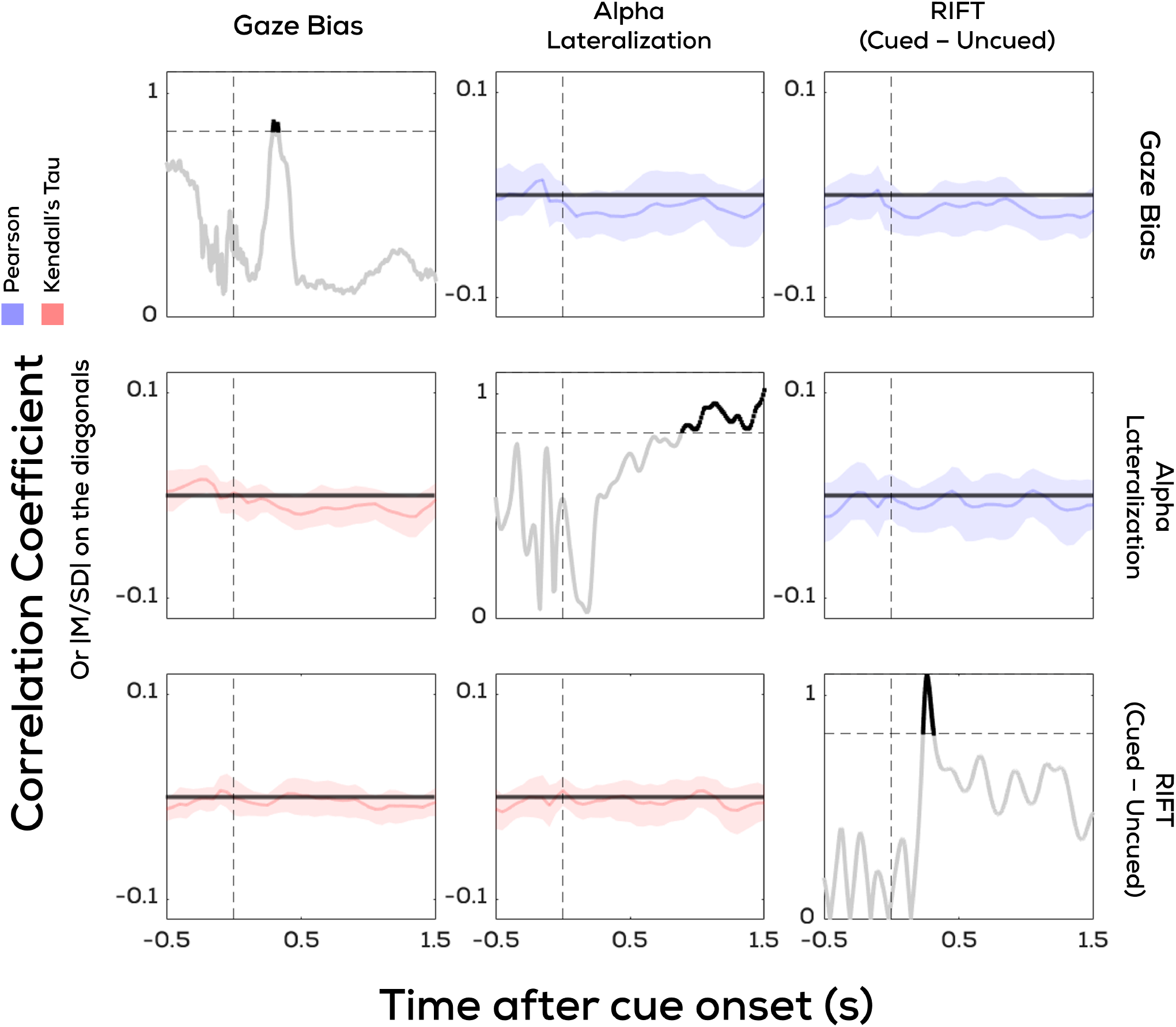
Proactive metrics do not correlate with each other. **Off-diagonal panels:** Timepoint-by-timepoint correlation coefficients of the gaze position bias, alpha lateralization, and RIFT response difference (between the cued and uncued locations) with each other. Shaded regions reflect boostrapped 95% CIs. RIFT modulations, alpha lateralization, and the gaze bias do not correlate with each other. **Diagonal panels:** Timepoint-by-timepoint mean/standard deviation of the gaze position bias, alpha lateralization, and RIFT attentional modulation. M/SD values in a selected range (0.85-1.1) are represented as thick sections, showing that all metrics here are comparable in their sensitivity with a peak M/SD of approximately 1.

Using a similar trial-wise approach, we also looked at whether the time-varying proactive measures (gaze position bias, alpha lateralization, RIFT responses from uncued and cued locations) correlated with time-invariant reactive measures tied to the target. RTs to the targets showed a negative correlation with the strength of the horizontal gaze bias, i.e., a stronger gaze bias predicted better performance (**Figure 7**). This effect was confirmed to be statistically significant for both parametric (p = 0.0025; cluster t-mass compared to bootstrapped distribution) and non-parametric (p = 0.006) tests using a cluster permutation test. This correlation was mainly localised to an early (0-500ms) time window. Beyond this, no correlations of any proactive measure were observed with target-evoked ERP lateralizations, or between RTs and alpha lateralizations or the RIFT modulation.

**Figure 7.**
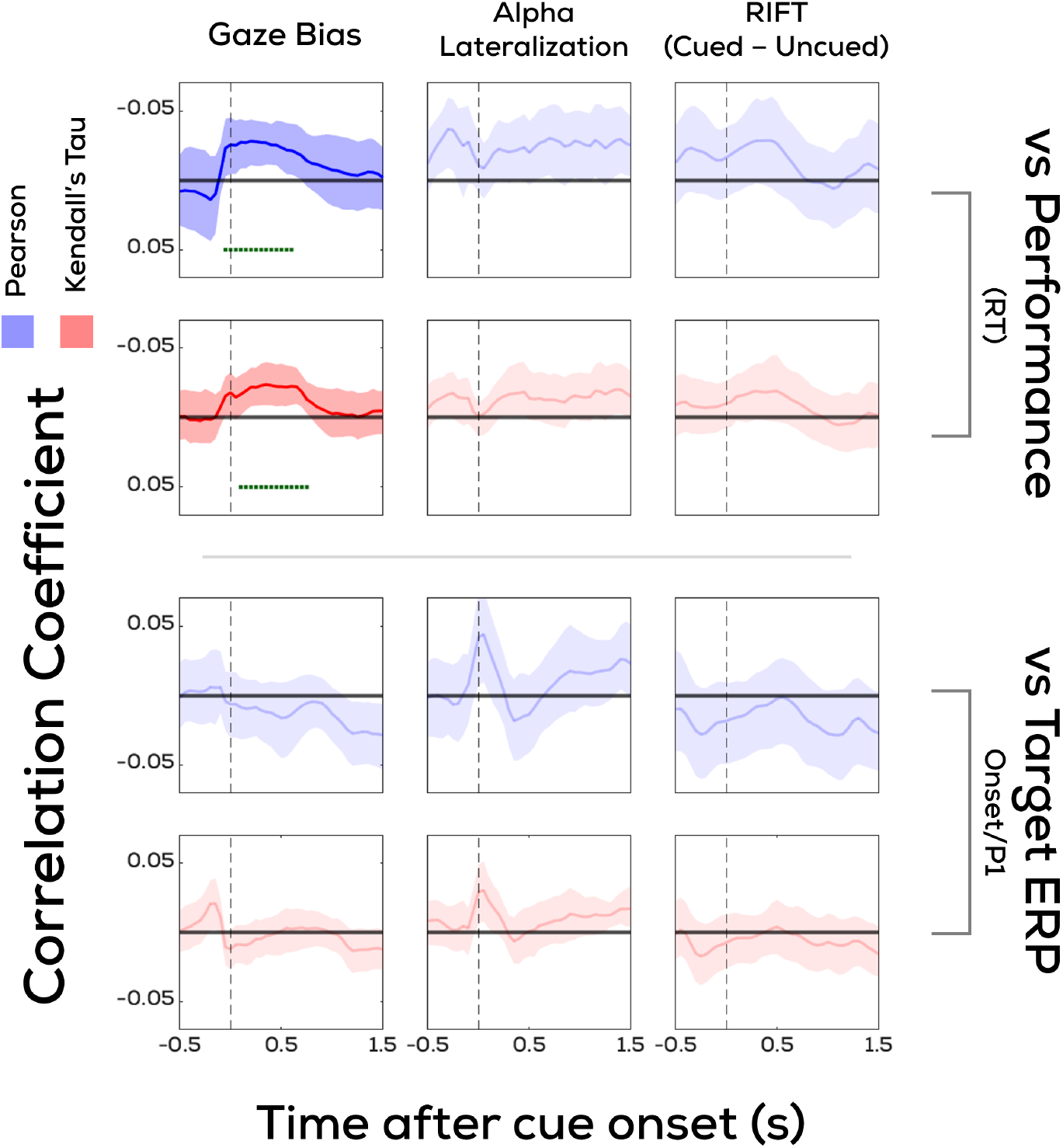
Of all shift-locked proactive measures, only gaze correlates with target reactions. Timepoint-by-timepoint correlation coefficients of the gaze position bias, alpha lateralization, and RIFT (difference in tagging amplitude between the cued and uncued regions) with response performance as measured by Reaction Time (RT), and lateralization of the target-Event Related Potential (ERP) response. Panels with significant correlations as measured by a cluster permutation test are shown in full opacity. Shaded regions reflect boostrapped 95% CIs. Note that the sign of the correlations with RTs are flipped such that upper values reflect better performance (lower RTs) and vice-versa. Thus, a stronger gaze bias correlates with better performance.

## Discussion

Covert attentional shifts and their various behavioral and neural correlates are widely studied across the domain of cognition. Despite the extensive literature surrounding each of these markers, the presence or absence of “attention” – as a single construct – at a particular location, or to a particular stimulus, is often concluded using only a subset, or even just one, of these methods. Given that attention is a complex, multi-level process, the underlying processes being measured by such attentional benefits may be very different depending on the marker used. However, the individual subprocesses that underlie shifts of spatial attention need not always be co-activated. Here, we demonstrate this: when measuring several common markers of attention in the same task that all reflect the attentional manipulation at the global level, the attentional benefits afforded by each of these do not all correlate with each other at a trialwise level.

First, we considered proactive measures of attention, i.e., those that attempt to directly measure a shift in the locus of attention before (or without) presenting visible stimuli at the attended/unattended locations. The proactive measures considered here (alpha oscillations, gaze position, and the RIFT response) all showed consistent attentional modulations, and we thus clearly measured a cue-induced shift of attention. These modulations, though statistically robust, did not correlate with one another. Thus, even within measures that all index the preparatory phase of attention, in a given trial the strength of one is not necessarily informative about the strength of the other. Though globally reflecting a shift of attention towards a cued location, these measures thus do not jointly read out the same underlying component of the attentional shifting process. One interpretation of this is that our proactive measures capture attentional subcomponents emerging at varying levels of the visual processing hierarchy. With RIFT, we measure retinotopic changes in excitability of the early visual cortex to luminance changes as attention is shifted (Drijvers et al., 2021; Duecker et al., 2025; Ferrante et al., 2023; Zhigalov et al., 2019). Alpha oscillations, though also a neural measure, are considered a separate gating mechanism that selectively controls the flow of information to downstream sections of the visual system (Jensen, 2024; Zhigalov & Jensen, 2020). Lastly, changes in gaze position are intrinsically a motor output. Thus, when studying attention, relying on one measure alone may lead to incorrect conclusions about the presence or absence of an attentional modulation.

We then posed the question: which of these independent processes link to our reactive measures (measures reflecting the differential response to targets at cued versus uncued locations – target ERPs and reaction times)? Only the cue-induced bias in gaze correlated with participant reports (specifically, to reaction time). We differentiate here between proactive and reactive measures as indices of two distinct steps of attentional deployment – the shift itself, and facilitation of stimulus processing, respectively – it would be natural to expect that both these steps occur together, since the former is ideally performed to facilitate the latter. That fact that gaze is the only proactive measure here that shows this link may relate to the processing levels at which the various measures operate. Gaze is similar to task performance in that it is a motor response; an outcome measure at the behavioural level. Any additional factors that may intervene between the neural response to a target stimulus and its report (e.g. decision-making, motor aspects) may similarly affect both the resulting shift of gaze position, as well as the report. This correlation is also in line with previous work showing that attentional cue-induced gaze biases index the quality of upcoming reports (Liu et al., 2022).

Though gaze does correlate with participant reports here, RIFT and alpha modulations do not. Why do these two measures so strongly reflect the spatial biasing of attention, if not to directly promote the processing of upcoming stimuli? By indexing neuronal excitability to visual changes at the attended location, and the strength with which this information is passed downstream, RIFT and alpha respectively provide information about the fidelity with which visual information from a given location is boosted as compared to other locations (Drijvers et al., 2021; Duecker et al., 2025; Ferrante et al., 2023; Jensen, 2024; Zhigalov & Jensen, 2020; Zhigalov et al., 2019). When RIFT and alpha are modulated by attention, a benefit to visual processing may be inferred, but this benefit may not always translate to a benefit in overt responses. Shifting attention often aims to enhance the perception of stimuli at attended locations in service of behavioural goals. However, the behavioural responses that we measure to target stimuli at these attended locations may not fully capture this enhancement, since those responses also reflect the consequences of other attentional subcomponents (eg. decision-making, motor aspects).

How does the independence of different markers of attention observed here relate to existing literature? Links between alpha oscillations and small eye movements have previously been investigated. Though correlations between microsaccadic eye movements and alpha lateralization have been observed, in the same studies it has also been shown that one can still be present if the other is absent (Liu et al., 2022, 2023). Similarly, it has been found that gaze can correlate with the N2pc ERP marker also considered here, but that the latter is not driven by the former (Liu et al., 2025). Rather than gaze biases and these other measures having a direct and causal relationship, they both thus likely reflect different underlying processes of attention that typically co-occur, but do not necessarily co-vary in magnitude, as we see here. The absence of any correlations between alpha oscillations and frequency tagging modulations has been reported several times (Antonov et al., 2020; Gundlach et al., 2020; Zhigalov & Jensen, 2020). Lastly, we have also found a similar absence of correlations between RIFT, alpha, and gaze biases in a previous study where we considered only proactive measures (Arora, Gayet, et al., 2025). Thus, there is support for our findings within existing literature.

Here, we include a relatively novel tool – RIFT – and show that neither attention-induced gaze biases nor alpha lateralization is correlated with the early visual sensitivity to luminance changes at the attended location as measured by RIFT. When inspecting the attentional modulations of these proactive measures, we observe a fairly sustained effect for RIFT and alpha lateralization (>1.5s), but a relatively short effect (*∼*200ms) for the gaze bias. This supports recent opinions in the field regarding microsaccadic eye movements as an index of shifting rather than sustaining covert attention (Van Ede, 2023; van Harmelen & van Ede, 2025), with RIFT and alpha lateralization then indexing the latter.

The absence of correlations seen here could result from our measures simply being too noisy to measure trialwise links rather than actually being uncorrelated. We address this in several ways. Firstly, we report robust attentional modulations, which would have been unlikely if our measures strongly reflected noise. Secondly, our reactive measures do correlate with each other as well as with the proactive gaze bias, confirming that these are sensitive enough to pick up on correlations. This of course does not eliminate the possibility that the gaze bias is the only proactive measure here sensitive enough to correlate with other measures. To address this, thirdly, we showed that SNR (as measured by M/SD) was roughly equivalent between all our proactive attentional modulations. We also conducted a more thorough subsampling analysis (Supplementary Figure S2), which showed that gaze is not the most sensitive of our measures. Rather, the attentional modulations of alpha lateralization were even more sensitive, indicating that the absence of other correlations is not driven by gaze biases being the most sensitive.

“Attention” is often the umbrella term to which findings about selective processing of visual stimuli are attributed. However, the various tools used to reveal these findings (participant reports, eye movements, alpha oscillations, and frequency tagging) capture distinct components associated with shifts of attention. We show this here by obtaining several such measures in the same attentional cueing task, and showing that they do not co-vary at the trialwise level. Attention research would therefore benefit from explicitly picking, and interpreting the findings of, the marker being used in the context of the underlying subprocess it measures. For example, when specifically concerned with cortical excitability in early visual cortex, RIFT would be appropriate. In summary, though all markers can pick up on attentional modulations, different markers of attention are not interchangeable: conclusions drawn about attention should be stated based on the attentional subcomponents that are reflected by the marker used.

## Data Availability Statement

Raw and processed EEG/Eyetracking data have been deposited in OSF and are publicly available at https://doi.org/10.17605/OSF.IO/CWR6Z.

## Author Contributions

Conceptualization, K.A., S.V.d.S., L.K., S.C., S.G..; data curation, K.A.; formal analysis, K.A., S.C.; funding acquisition, S.V.d.S.; investigation, K.A.; methodology, K.A., S.C., and S.G.; project administration, K.A., S.C.; resources, S.V.d.S.; software, K.A., S.C.; supervision, S.V.d.S., L.K., S.C., S.G.; visualization, K.A.; writing—original draft preparation, K.A., S.C., and S.G.; writing—review and editing, K.A., S.V.d.S., L.K., S.C., S.G.

## Supplementary Figures

**Figure S1.**
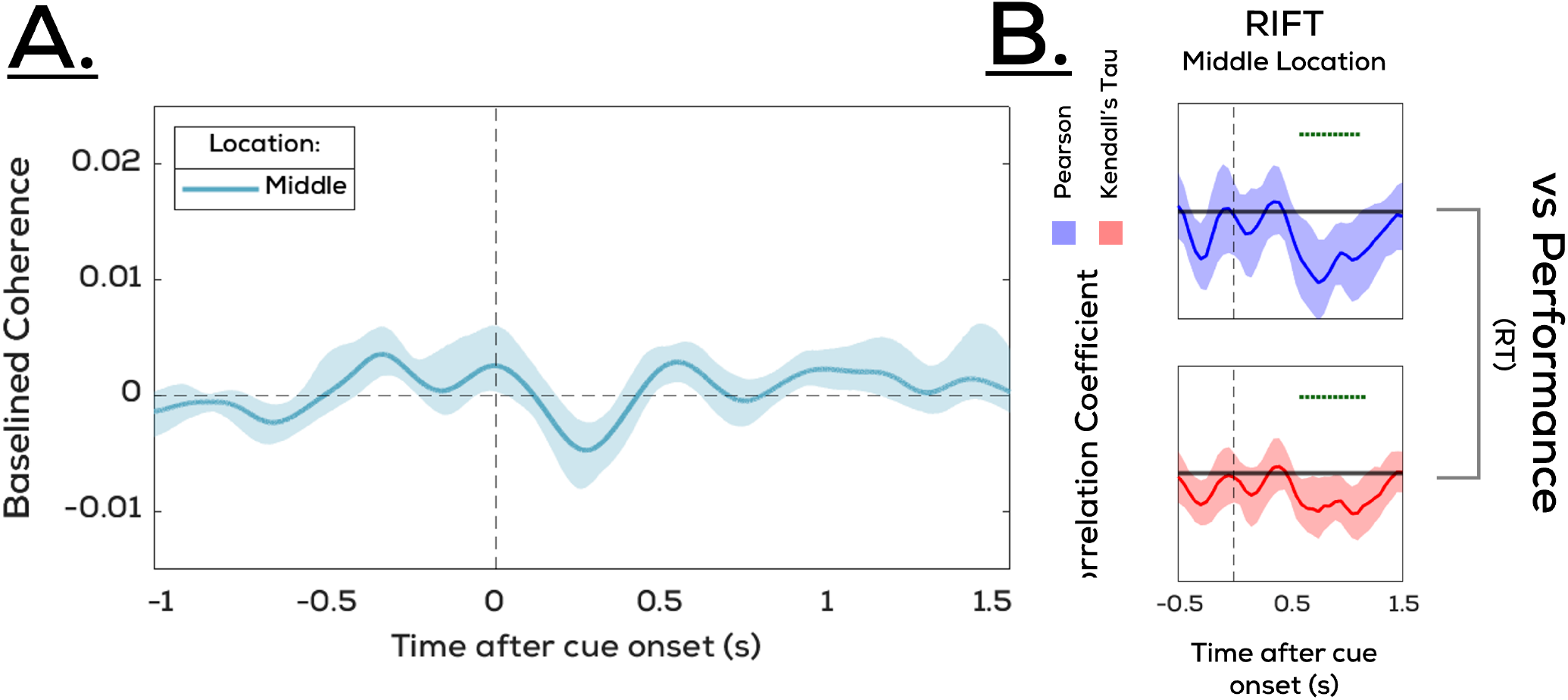
Tagging below fixation (middle location) suggests inability to sustain covert attention. In addition to the “cued” and “uncued” RIFT tags discussed in the results of the main manuscript, we also tagged a square directly below fixation, i.e., the “middle” location. This was not considered in the main analysis since the attentional modulations resulting from cue-induced modulations in the RIFT response require only the difference between the tagging responses of the cued and uncued locations. **A**. However, we also observed a trend in the middle location response following the cue. Though not statistically significant, the RIFT response from the middle location was relatively lower than the baseline for a brief moment, before returning to baseline levels around 500ms after the cue. Secondly, we observed that unlike the cued vs. uncued RIFT response differences, **B**. the middle location RIFT response in fact showed a correlation with participant reports (RT), with a stronger RIFT response from the middle location correlating with worse performance. However, compared to the gaze correlations this occurred later after the cue (500-1000ms as opposed to 0-500ms). Together, these findings suggest the following possibility: participants did not always successfully sustain attention at the cued location until the next cue occured. Instead, on a subset of trials, attention moved to fixation (i.e., in line with gaze position). On these trials, a higher RIFT response would be observed at the middle location in the later intervals following cue onset, and this attentional shift to fixation should result in poorer performance on targets appearing at the cued location. Though this does not have major implications for our task where we look at shifts of attention, this may imply that participants find it difficult to sustain covert attention peripherally even at the cost of performance.

**Figure S2.**
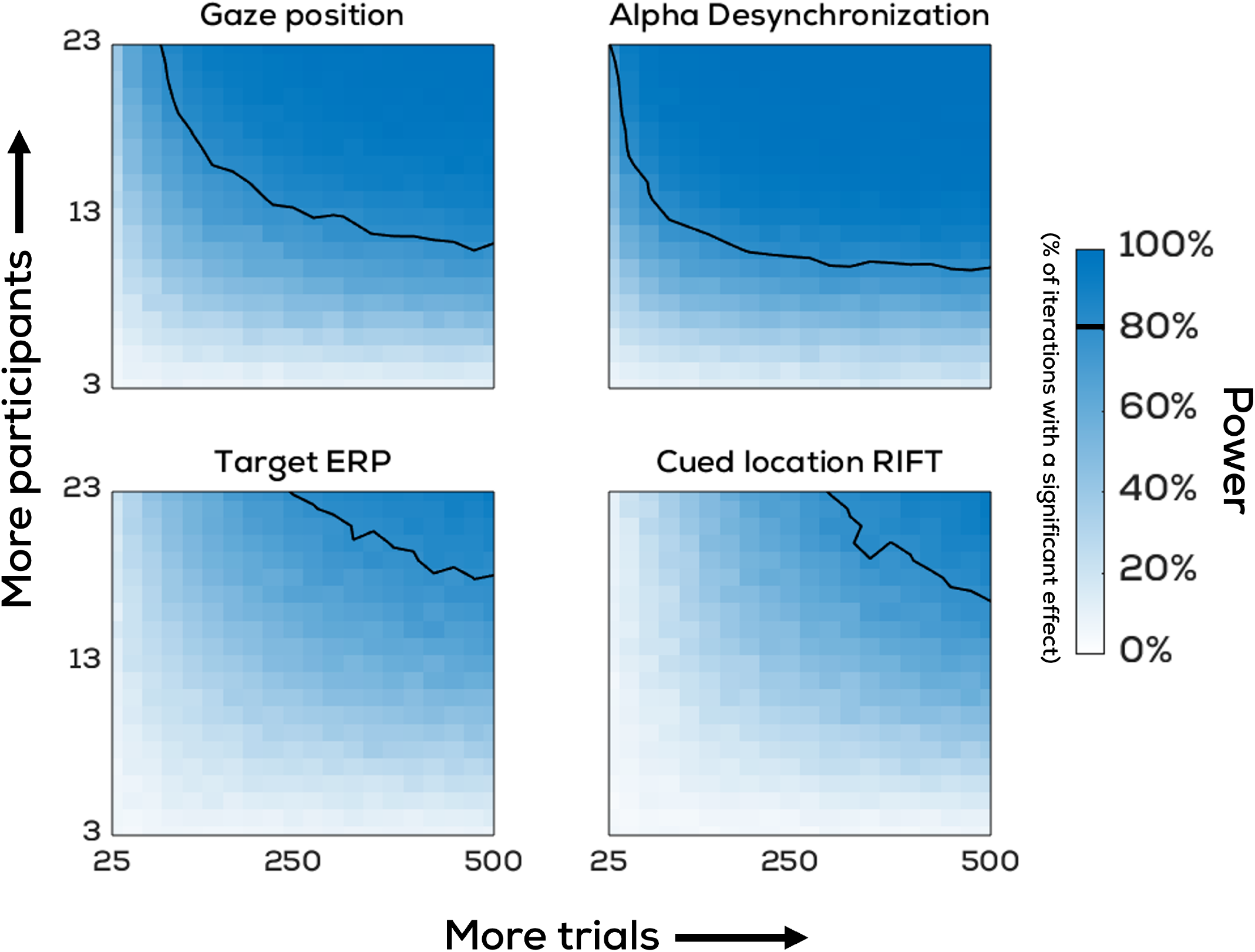
Subsampling analyses allow for qualitative sensitivity comparisons across measures. **Background:** Having several common measures of attentional shifts in the same dataset, all of which showed reliable attentional modulations, we ran an additional analysis to determine whether the strength of these modulations would have been retained given a number of trials and participants smaller than that of the full dataset (23 participants, 700 trials per participant). For sub-samplings of n participants (ranging from 3-23) and m trials (ranging from 25-500 in increments of 25), we ran 1000 iterations each where we randomly sampled m trials from n participants with replacement. For each of these 1000 subsets, a participant-wise time-averaged attentional modulation of alpha lateralization, gaze position bias, target-locked ERPs, and RIFT responses from the cued location, was obtained using the corresponding time intervals significant at the group level. Lastly, we then determined the percentage of iterations for which a significant effect was observed in each of these metrics (i.e., statistical power). **Figure details:** Proportion of iterations (out of 1000) where randomly sampled subsets of a fixed number of trials (horizontal axis) and participants (vertical axis) showed a significant attentional modulation for gaze position, alpha lateralization, target-evoked ERP, and RIFT responses from the cued location. Solid contour line reflects 80% power threshold.

## References

Antonov, P. A., Chakravarthi, R., & Andersen, S. K. (2020). Too little, too late, and in the wrong place: Alpha band activity does not reflect an active mechanism of selective attention [Publisher: Elsevier]. NeuroImage, 219, 117006. 10.1016/j.neuroimage.2020.117006

Arora, K., Gayet, S., Kenemans, J. L., Van der Stigchel, S., & Chota, S. (2025). Dissociating external and internal attentional selection [Publisher: Elsevier]. iScience, 28(4). 10.1016/j.isci.2025.112282

Arora, K., Husta, C., Bouwkamp, F., Seijdel, N., Bai, S., Han, Q., Kenemans, L., Van Der Stigchel, S., Gayet, S., Spaak, E., Chota, S., & Drijvers, L. (2025, October 28). A collaborative guide to rapid invisible frequency tagging (RIFT): Methods, insights, and recommendations. 10.31234/osf.io/edshx_v1

Carrasco, M., Penpeci-Talgar, C., & Eckstein, M. (2000). Spatial covert attention increases contrast sensitivity across the CSF: Support for signal enhancement [Publisher: Elsevier]. Vision research, 40(10), 1203– 1215. 10.1016/S0042-6989(00)00024-9

Dietz, L., Strauch, C., Arora, K., Van der Stigchel, S., Chota, S., & Gayet, S. (2025). Anticipated relevance prepares visual processing for efficient memory-guided selection [Publisher: Cold Spring Harbor Laboratory]. bioRxiv, 2025–08. 10.1101/2025.08.20.671232

Doherty, J. R., Rao, A., Mesulam, M. M., & Nobre, A. C. (2005). Synergistic effect of combined temporal and spatial expectations on visual attention [Publisher: Society for Neuroscience]. Journal of Neuroscience, 25(36), 8259–8266. 10.1523/JNEUROSCI.1821-05.2005

Drijvers, L., Jensen, O., & Spaak, E. (2021). Rapid invisible frequency tagging reveals nonlinear integration of auditory and visual information. Human Brain Mapping, 42(4), 1138–1152. 10.1002/hbm.25282

Duecker, K., Shapiro, K. L., Hanslmayr, S., Griffiths, B. J., Pan, Y., Wolfe, J. M., & Jensen, O. (2025). Guided visual search is associated with target boosting and distractor suppression in early visual cortex [Publisher: Nature Publishing Group]. Communications Biology, 8(1), 1–11. 10.1038/s42003-025-08321-3

Engbert, R., & Kliegl, R. (2003). Microsaccades uncover the orientation of covert attention [Publisher: Elsevier]. Vision research, 43(9), 1035–1045. 10.1016/S0042-6989(03)00084-1

Farell, B., & Pelli, D. G. (1999). Psychophysical methods, or how to measure a threshold and why [Publisher: Oxford University Press New York, NY]. Vision research: A practical guide to laboratory methods, 5, 129–136. https://doi.org/dx.doi.org/10.1093/acprof:oso/9780198523192.003.0005

Ferrante, O., Zhigalov, A., Hickey, C., & Jensen, O. (2023). Statistical learning of distractor suppression downregulates prestimulus neural excitability in early visual cortex [Publisher: Society for Neuroscience]. Journal of Neuroscience, 43(12), 2190–2198. 10.1523/JNEUROSCI.1703-22.2022

Foster, J. J., Sutterer, D. W., Serences, J. T., Vogel, E. K., & Awh, E. (2017). Alpha-band oscillations enable spatially and temporally resolved tracking of covert spatial attention. Psychological Science, 28(7), 929–941. 10.1177/0956797617699167

Foxe, J. J., Simpson, G. V., & Ahlfors, S. P. (1998). Parietooccipital 1 0hz activity reflects anticipatory state of visual attention mechanisms [Publisher: LWW]. Neuroreport, 9(17), 3929–3933. 10.1097/00001756-199812010-00030

Gould, I. C., Rushworth, M. F., & Nobre, A. C. (2011). Indexing the graded allocation of visuospatial attention using anticipatory alpha oscillations. Journal of Neurophysiology, 105(3), 1318–1326. 10.1152/jn.00653.2010

Gundlach, C., Moratti, S., Forschack, N., & Müller, M. M. (2020). Spatial attentional selection modulates early visual stimulus processing independently of visual alpha modulations [Publisher: Oxford University Press]. Cerebral Cortex, 30(6), 3686–3703. 10.1093/cercor/bhz335

Hafed, Z. M., & Clark, J. J. (2002). Microsaccades as an overt measure of covert attention shifts [Publisher: Elsevier]. Vision research, 42(22), 2533–2545. 10.1016/S0042-6989(02)00263-8

Hershman, R., Henik, A., & Cohen, N. (2018). A novel blink detection method based on pupillometry noise. Behavior Research Methods, 50(1), 107–114. 10.3758/s13428-017-1008-1

Hillyard, S. A., & Münte, T. F. (1984). Selective attention to color and location: An analysis with event-related brain potentials. Perception & Psychophysics, 36(2), 185–198. 10.3758/BF03202679

Hustá, C., Meyer, A., & Drijvers, L. (2025). Using rapid invisible frequency tagging (RIFT) to probe the neural interaction between representations of speech planning and comprehension [Publisher: MIT Press 255 Main Street, 9th Floor, Cambridge, Massachusetts 02142, USA …]. Neurobiology of Language, 1–36. http://dx.doi.org/10.1162/nol_a_00171

Jensen, O. (2024). Distractor inhibition by alpha oscillations is controlled by an indirect mechanism governed by goal-relevant information [Publisher: Nature Publishing Group UK London]. Communications Psychology, 2(1), 36. 10.1038/s44271-024-00081-w

Kleiner, M., Brainard, D., & Pelli, D. (2007). What’s new in psychtoolbox-3? [Publisher: Pion Ltd.].

Liu, B., Kong, S., & van Ede, F. (2025). Microsaccades strongly modulate but do not directly cause the EEG n2pc marker of spatial attention [Publisher: Public Library of Science San Francisco, CA USA]. Plos Biology, 23(9), e3003418. 10.1371/journal.pbio.3003418

Liu, B., Nobre, A. C., & van Ede, F. (2022). Functional but not obligatory link between microsaccades and neural modulation by covert spatial attention [Publisher: Nature Publishing Group UK London]. Nature Communications, 13(1), 3503. http://dx.doi.org/10.1038/s41467-022-31217-3

Liu, B., Nobre, A. C., & van Ede, F. (2023). Microsaccades transiently lateralise EEG alpha activity [Publisher: Elsevier]. Progress in Neurobiology, 224, 102433. 10.1016/j.pneurobio.2023.102433

Luck, S. J., & Kappenman, E. S. (2012). Electrophysiological correlates of the focusing of attention within complex visual scenes: N2pc and related ERP components. The Oxford handbook of event-related potential components, 329–360.

Luck, S. J., Woodman, G. F., & Vogel, E. K. (2000). Eventrelated potential studies of attention [Publisher: Elsevier]. Trends in cognitive sciences, 4(11), 432– 440. 10.1016/S1364-6613(00)01545-X

Mangun, G. R., & Hillyard, S. A. (1988). Spatial gradients of visual attention: Behavioral and electrophysiological evidence [Publisher: Elsevier]. Electroencephalography and clinical Neurophysiology, 70(5), 417–428.

Maris, E., & Oostenveld, R. (2007). Nonparametric statistical testing of EEG-and MEG-data [Publisher: Elsevier]. Journal of neuroscience methods, 164(1), 177–190.

Minarik, T., Berger, B., & Jensen, O. (2023). Optimal parameters for rapid (invisible) frequency tagging using MEG [Publisher: Elsevier]. NeuroImage, 281, 120389. 10.1016/j.neuroimage.2023.120389

Morgan, M., Ward, R., & Castet, E. (1998). Visual search for a tilted target: Tests of spatial uncertainty models. The Quarterly Journal of Experimental Psychology Section A, 51(2), 347–370. 10.1080/713755766

Müller, M. M., Malinowski, P., Gruber, T., & Hillyard, S. A. (2003). Sustained division of the attentional spotlight [Publisher: Nature Publishing Group UK London]. Nature, 424(6946), 309–312. 10.1038/nature01812

Müller, M. M., Picton, T. W., Valdes-Sosa, P., Riera, J., Teder-Sälejärvi, W. A., & Hillyard, S. A. (1998). Effects of spatial selective attention on the steady-state visual evoked potential in the 20–28 hz range [Publisher: Elsevier]. Cognitive brain research, 6(4), 249– 261. 10.1016/S0926-6410(97)00036-0

Neville, H. J., & Lawson, D. (1987). Attention to central and peripheral visual space in a movement detection task: An event-related potential and behavioral study. II. congenitally deaf adults [Publisher: lsevier]. Brain research, 405(2), 268–283. http://dx.doi.org/10.1016/0006-8993(87)90296-4

Norcia, A. M., Appelbaum, L. G., Ales, J. M., Cottereau, B. R., & Rossion, B. (2015). The steady-state visual evoked potential in vision research: A review [Publisher: The Association for Research in Vision and Ophthalmology]. Journal of vision, 15(6), 4–4. 10.1167/15.6.4

Oostenveld, R., Fries, P., Maris, E., & Schoffelen, J.-M. (2011). FieldTrip: Open source software for advanced analysis of MEG, EEG, and invasive electrophysiological data. Computational Intelligence and Neuroscience, 2011, 1–9. 10.1155/2011/156869

Posner, M. I. (1980). Orienting of attention. Quarterly Journal of Experimental Psychology, 32(1), 3–25. 10.1080/00335558008248231

Rugg, M. D., Milner, A. D., Lines, C. R., & Phalp, R. (1987). Modulation of visual event-related potentials by spatial and non-spatial visual selective attention [Publisher: Elsevier]. Neuropsychologia, 25(1), 85– 96. 10.1016/0028-3932(87)90045-5

Sauseng, P., Klimesch, W., Stadler, W., Schabus, M., Doppelmayr, M., Hanslmayr, S., Gruber, W. R., & Birbaumer, N. (2005). A shift of visual spatial attention is selectively associated with human EEG alpha activity. European Journal of Neuroscience, 22(11), 2917– 2926. 10.1111/j.1460-9568.2005.04482.x

Seijdel, N., Marshall, T. R., & Drijvers, L. (2023). Rapid invisible frequency tagging (RIFT): A promising technique to study neural and cognitive processing using naturalistic paradigms [Publisher: Oxford University Press]. Cerebral Cortex, 33(5), 1626–1629.

Serences, J. T., & Kastner, S. (2014). A multi-level account of selective attention [Publisher: Oxford University Press Oxford]. The Oxford handbook of attention, 76–104.

Stoletniy, A. S., Alekseeva, D. S., Babenko, V. V., Anokhina, P. V., & Yavna, D. V. (2022). The n2pc component in studies of visual attention. Neuroscience and Behavioral Physiology, 52(8), 1299–1309. 10.1007/s11055-023-01359-y

Van Ede, F. (2023). Do microsaccades track shifting but not sustaining covert attention? Proceedings of the National Academy of Sciences, 120(35), e2309431120. 10.1073/pnas.2309431120

Van Ede, F., Chekroud, S. R., & Nobre, A. C. (2019). Human gaze tracks attentional focusing in memorized visual space [Publisher: Nature Publishing Group UK London]. Nature human behaviour, 3(5), 462–470. 10.1038/s41562-019-0549-y

van Harmelen, A. M., & van Ede, F. (2025). Microsaccades track shifting but not necessarily maintaining covert visual-spatial attention [Publisher: Cold Spring Harbor Laboratory]. bioRxiv, 2025–07.

Wang, D., Arora, K., Theeuwes, J., Stigchel, S. V. d., Gayet, S., & Chota, S. (2025). Dynamic competition between bottom-up saliency and top-down goals in early visual cortex [Publisher: Cold Spring Harbor Laboratory]. bioRxiv, 2025–08. 10.1101/2025.08.22.671530

Zhigalov, A., Herring, J. D., Herpers, J., Bergmann, T. O., & Jensen, O. (2019). Probing cortical excitability using rapid frequency tagging [Publisher: Elsevier]. NeuroImage, 195, 59–66.

Zhigalov, A., & Jensen, O. (2020). Alpha oscillations do not implement gain control in early visual cortex but rather gating in parieto-occipital regions [_eprint: https://onlinelibrary.wiley.com/doi/pdf/10.1002/hbm.25 Human Brain Mapping, 41(18), p5176–5186. 10.1002/hbm.25183

